# Stress responses and dynamic equilibrium: Key determinants of aging in the *C. elegans clk-1* mutant

**DOI:** 10.1101/2024.08.21.609027

**Authors:** Jose Carracedo-Gonzalez, Fausto Arellano-Carbajal, Etzel Garrido, Roberto Alvarez-Martinez

## Abstract

Systems biology offers valuable insights into aging by integrating experimental data with mathematical models and bioinformatics tools. Long-lived mutants of *C. elegans*, particularly *clk-1*, have provided extensive data on the aging mechanisms. The *clk-1* gene, which encodes a ubiquitin precursor, shows a pleiotropic phenotype characterized by slow behavior, high mitochondrial ROS levels, autophagy, and metabolic changes. However, the link between these changes and lifespan extension remains unclear. Using a Boolean network, we modeled genetic interactions and derived differential equations for a continuous approach. Our results highlight that *aak-2* (AMPK) is crucial for *clk-1* lifespan extension owing to its role in stress response regulation. We introduced a health index based on the attrition of neuromuscular behaviors to assess the health of various strains. Our findings suggest that while stress responses may enhance lifespan, overall health is determined by the extent of the damage.

## Introduction

The nature of the aging process has captivated scientists for decades, yet its underlying mechanisms and primary drivers remain only partially understood. To elucidate these complexities, researchers have extensively employed model organisms, with the nematode *Caenorhabditis elegans* proving to be particularly useful. This worm has a mean lifespan of merely 20 days and approximately 80% of its genes have human orthologous genes (1) thus offering a unique window into the genetic underpinnings of aging. Particularly, mutant strains of this organism have proven invaluable for exploring the genetic modulation of lifespan. Investigations have spotlighted gene deletions that prolong life, implicating pathways such as insulin signaling (2), dietary restriction mechanisms (3), and mitochondrial dynamics (4). In particular, *clk-1* gene, a precursor for ubiquinone synthesis, has garnered attention due to its mutation’s extensive impact on both lifespan and behavioral phenotype, provoking pleiotropic effects that transcend simple metabolic adjustments (5).

Over the past quarter-century, theories attempting to elucidate the lifespan extension observed in *clk-1* mutants have ranged from the metabolic rate hypothesis (6) to propositions of singular regulatory processes (7,8) and the more nuanced hormesis theory (9). Each theory contributes to our understanding yet underscores the complexity of the aging process, highlighting the need for a holistic approach to deciphering the multitude of molecular interactions and changes.

In addressing this complexity, logical modeling, mainly through constructing Boolean networks, emerges as a powerful tool. Capable of integrating diverse types of interactions, simulating system dynamics, and identifying stable states, Boolean or logical networks have been used extensively in various biological systems since their inception (10). Notable applications include replicating T cell differentiation (11), exploring mutant phenotypes in *Arabidopsis thaliana* (12), and evaluating pharmacological interventions (13,14). This modeling approach holds particular promise for unraveling the *clk-1* mutant paradox, where reduced ubiquinone levels and resultant ROS accumulation paradoxically extend lifespan, suggesting a complex interplay of molecular interactions awaiting integration into a comprehensive model.

A unique advantage of logical modeling is its ability to simulate perturbations, mirroring mutant conditions such as the *clk-1;aak-2* double mutant, which reverses the lifespan extension of *clk-1*. This mutation, affecting the AMPK kinase alpha subunit, modifies behaviors like the pharyngeal pumping rate, offering insights into the subtle relationship between genetic mutations, lifespan, and healthspan (15). While lifespan extension in mutants is noteworthy, alterations in behavior highlight the relevance of healthspan — the period of life spent in good health — which remains challenging to quantify due to its subjective nature and the difficulty in pinpointing the transition to functional decline (16, 17).

Here, we adopt a systems biology approach to dissect the phenotypic outcomes of single-gene mutations in *clk-1* and the *clk-1*;*aak-2* double mutant through logical modeling. By analyzing discrete and continuous system dynamics to identify attractors and correlating these with measured behaviors such as pharyngeal pumping, swimming, and defecation, we aim to establish a comprehensive health condition profile for these strains by proposing a novel health index estimation based on the overall decline in behaviors over nine days. Overall, this study aims to understand the aging process by creating a discrete Boolean network grounded in three decades of literature. The network’s steady states were analyzed using both discrete and continuous approaches, incorporating expression level data to validate the model. This quantitative approach helped reveal the relationship between the model and in vivo data, highlighting the role of stress response and the dynamic nature of aging.

## Materials and methods

### Construction and analysis of the *clk-1* Regulatory Network Model

To build a comprehensive model for the *clk-1* regulatory network we thoroughly researched the existing literature to gather all documented interactions relevant to *clk-1*. We were able to include the complete set of interactions to ensure that our regulatory network accurately reflects the current understanding of this organism model. In the network, each node symbolizes a distinct molecular component, such as genes, proteins, or small molecules, integral to the *clk-1* network. The edges that connect these nodes denote the regulatory influence — positive or negative — one component exerts on another.

We used the Cytoscape software (18) to visualize the biological network depicted in the graph. Through Cytoscape, we represented the *clk-1* regulatory graph in a user-friendly format, facilitating a simplified exploration of its structure and dynamics. Furthermore, we enhanced the precision and predictive capability of the model using the R-package BoolNet (19). This package was the computational framework for simulating the *clk-1* network and identifying its stable states. These are crucial in representing network configurations that may endure over time, potentially correlating with specific phenotypes.

Additionally, we utilized the R-package BoolNetPerturb (20) to simulate the effects of various perturbations on the *clk-1* network, gaining valuable insights into its resilience and adaptability. Furthermore, the deSolve R package (21) was used to delve into the network’s continuous dynamic behaviors, leading to a more comprehensive understanding of its temporal dynamics under different conditions.

### *C. elegans* strains and culture

Nematodes were cultured in standard conditions on Nematode Growth Medium (NGM) plates and fed with the *Escherichia coli* strain OP50 at 20°C temperature (22). In the experiments, Bristol N2 was used as the wild-type strain while the mutant strains used were *aak-2(ok524)* and *clk-1(qm30)*. These strains were provided by the Caenorhabditis Genetics Center (CGC), which is funded by the NIH Office of Research Infrastructure Programs (P40 OD010440). To create a double mutant, *aak-2(ok524)* males were crossed with *clk-1(qm30)* hermaphrodites. F2 progeny were then separated on individual plates to identify the double mutants using the polymerase chain reaction (PCR) technique. To identify the *aak-2* mutation, the following primer sequences were utilized: forward TTCCTGGCAACACCATAAGC and reverse CTCCAGAAAGTCTGGAGTTG. On the other hand, to detect the *clk-1* mutation, the primer sequences used were forward TGTCGGTTCAGCACTTCTAG and reverse AGTATTGTCCGTGTCAGGAC. This method allowed for precise identification of the desired genetic configurations.

### Measurement of Neuromuscular Behaviors

Our study assessed neuromuscular behaviors using the methodologies outlined by Hart (23), focusing on a trio of specific behaviors: pharyngeal pumping, swimming, and defecation. These behaviors were selected as indicators of neuromuscular activity and coordination, offering insights into the nematodes’ physiological health and functional status under investigation.

The experimental protocol started with nematodes that had reached one day of adulthood. A cohort of 20 worms was selected to ensure age and developmental stage uniformity. This selection criterion was crucial for minimizing variability in neuromuscular function that might arise from differences in maturation. The behaviors of these worms were then systematically observed and recorded over nine days, with assessments conducted at 48-hour intervals. This longitudinal approach allowed us to capture possible changes in neuromuscular activity over time, providing a comprehensive overview of each behavior.

We measured pharyngeal pumping and swimming behaviors, involving three replicates for each worm at every observation point. Pharyngeal pumping, a critical indicator of feeding behavior and metabolic activity, was quantified by counting the number of pharyngeal contractions within a predefined time frame (30 seconds). Swimming behavior, reflecting motor coordination and muscle function, was evaluated based on the worms’ frequency of tail waving in an aquatic environment, within a predefined time frame (60 seconds). The defecation behavior of each worm was measured with a single replicate, we recorded the time between defecation cycles, characterized by periodic expulsion events, within a 10 minutes time frame, to gain insight into the regulation of the gastrointestinal and neuromuscular systems.

### Survival analysis

Our survival analysis protocol was designed following the procedures delineated by Park et al. (24) to ensure the reliability and reproducibility of our results. This approach allowed for the precise monitoring of nematode longevity under controlled conditions. The experimental setup began with selecting nematodes at the L4 larval stage, an easily identifiable stage in their development. These selected individuals were then carefully transferred onto fresh Nematode Growth Medium (NGM) plates. This transfer was performed to standardize the starting conditions for each nematode, thereby reducing variability in survival outcomes that might arise from differences in environmental factors or developmental stages at the onset of the assay.

Twenty-four hours post-transfer, marking the transition of the nematodes to one-day adults, the survival assays officially commenced, with this time point designated as t = 1. Starting the assays at the adult stage ensured that the survival data captured pertained exclusively to the adult phase of the nematode life cycle, facilitating the analysis on adult longevity.

Survival was assessed every 24 hours. Nematodes that deviated from the experimental parameters, such as those that crawled off the plate, exhibited bagging (a condition where eggs hatch inside the mother, leading to her death), or exploded, were censored at the time these events were observed. The censoring of these individuals was critical for the integrity of the survival analysis, as it accounted for non-standard mortality causes that do not directly relate to the natural lifespan of the organism. The nematodes were transferred to new NGM plates every 48 hours to maintain optimal conditions and minimize confounding factors such as overcrowding or depletion of food resources. This bi-daily transfer protocol was pivotal in ensuring that environmental degradation did not influence the observed survival outcomes over time. Additionally, this strategy prevented the inadvertent inclusion of new adults from potential reproduction, thus maintaining the purity of the adult cohort under study.

### Health index estimation

In our study, we developed a nuanced approach to estimate the health index of nematodes, leveraging the variables derived from our behavioral analyses. This estimation process was designed to encompass individual and aggregated behavioral metrics, providing a holistic view of the nematodes’ physiological state over time.

### Aggregation of Behavioral Data into a Health Index

To construct the health index we first relativize all the behavioral data by converting it to a relative scale. This standardization facilitated the comparison and aggregation of diverse behaviors on a standard metric, enhancing the interpretability of our analysis. Data from different behaviors were cumulatively added for the aggregation process to form a composite score. Notably, in the case of defecation—a behavior that increases its length over time—its relative metric was subtracted from the composite of the other behaviors. This adjustment was crucial for accurately reflecting the contribution of each behavior to the overall health index, acknowledging that not all behaviors equally signify healthiness.

### Symbolic Regression for Health Index Estimation

The initial method used for estimating the health index was symbolic regression, executed through the software Eureqa (25). Symbolic regression diverges from traditional regression by searching for the mathematical expression that best fits the relationship within the data rather than fitting the data to a predefined model. Through this process, we first identified the optimal function that exhibited the highest fidelity to the behavioral data patterns and then we calculated its definite integral over the observed period. This integral, representing the area under the curve generated by the function, served as a quantitative measure of the cumulative health index across the lifespan of the nematodes.

### Direct Area Calculation from Behavioral Data

Additionally to the symbolic regression approach, we employed a direct method to estimate the health index by calculating the area under the data curve. This method drew lines through the behavioral data’s first, second, or third quartiles for each measurement day. By connecting these points across the observation period, we delineated polygons whose boundaries were defined by the quartile lines and the temporal axis representing the consecutive days. The area enclosed by these polygons was then computed, offering an alternative measure of the health index that is directly derived from the raw behavioral data.

### Statistical analysis

In our study, the statistical analysis assessed the effects of genetic variation and temporal dynamics on neuromuscular behaviors and survival across different nematode strains. For the variables of: pharyngeal pumping frequency, defecation cycle speed and waving frequency during swimming. We performed a two-way mixed Analysis of Variance (ANOVA) with nematode strain (fixed factor), time (as random factor) and their interaction as sources of variation. This analysis facilitated the discernment of strain-specific behavioral profiles and their progression over time, allowing us to pinpoint any significant deviations attributable to genetic differences or temporal changes. The test allows us to evaluate the interaction effect between strain and time, providing insights into whether the impact of genetic variation on neuromuscular behavior changes over the lifespan of the nematodes.

The Cox proportional hazards regression model was employed to analyze survival data. The Cox regression analysis was performed to verify the proportionality of the hazard assumption, employing both graphical diagnostics and statistical tests. This step ensured the validity of the model’s findings. The survival curves generated from this model were then statistically compared, with the resulting p-values providing a basis for assessing the significance of differences observed between strains, all statistical analyses were conducted using the R programming language (26)

## Results

### The Boolean network integrates all the known interactions for the *clk-1* mutant

The resulting Boolean network model, depicted in Figure 1, incorporates 29 distinct molecular components and 46 specific interactions. These components span a variety of biological entities, including individual proteins and entire metabolic pathways, illustrating the diverse nature of the *clk-1* regulatory network (as detailed in Table 1). The model depicts key processes such as mitochondrial unfolded protein response (UPRmt), autophagy, mitochondrial fusion, and lipid metabolism, emphasizing the wide range of cellular mechanisms influenced by *clk-1* mutations.

**Table 1:**
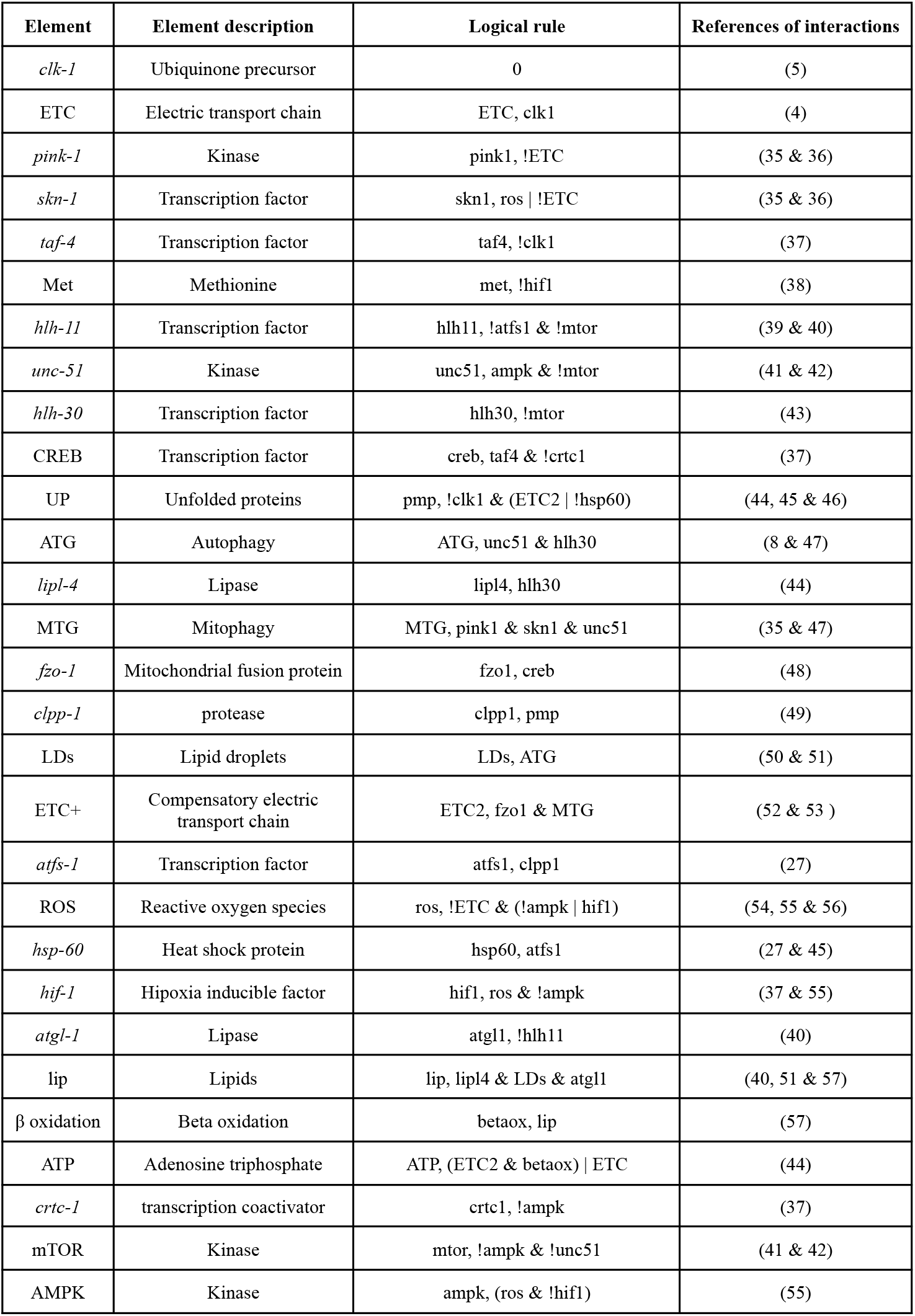
Network components and their interactions. Logical rules are defined in boolnet nomenclature.

**Figure 1.**
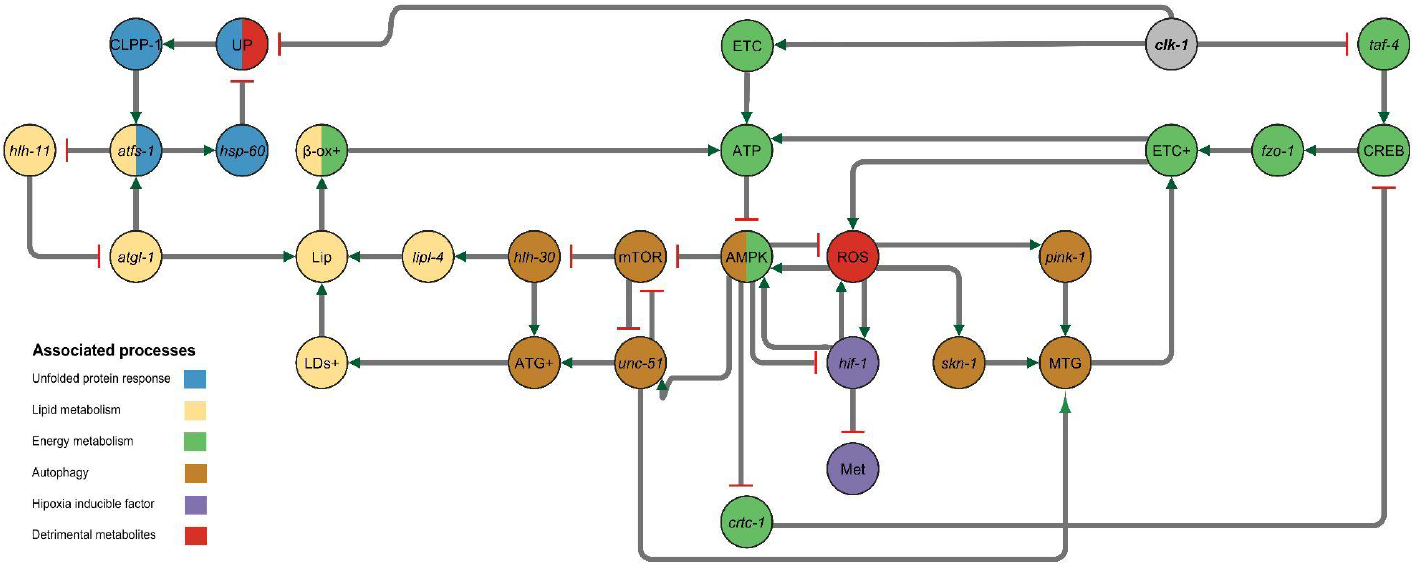
Logical regulatory network of *clk-1* mutant molecular interactions. The model was designed to study the regulatory landscape in the *C. elegans clk-1* mutant. The nodes are divided into six categories: UPRmt nodes, lipid metabolism nodes, energy metabolism nodes, autophagy nodes, hypoxia response nodes, and detrimental metabolites nodes. A color code denotes these categories. The arcs connecting the nodes represent positive (green arrow) or negative (red line) regulatory interactions.

Based on empirical evidence, each interaction was encoded using logical rules to translate the *clk-1* interactions into a functional model (referenced in Table 1). This approach enabled the computational simulation of the network’s behavior, providing insights into the dynamic interplay among the components. A particularly complex aspect of the model is the feedback loop involving AMP-activated protein kinase (AMPK), reactive oxygen species (ROS), and hypoxia-inducible factor 1 (HIF-1). The density of interactions within this loop posits significant challenges for simulating the network’s dynamics, requiring the adoption of a discrete synchronous update model to manage the computational complexity.

Given the complexity observed in the feedback loop among AMPK, ROS, and HIF-1, we opted to implement two distinct configurations to accurately model the interactions between these critical elements. The first configuration adopts a simplified approach, focusing on the minimal interactions required to activate or inhibit the elements. This configuration was designed to reduce the model’s complexity, enabling a clearer understanding of the fundamental mechanisms (Figure 2 A and C). The other configuration contemplates the whole interactions as described in the literature; this is biologically more accurate, but it is not feasible for the discrete model, due to the constant positive and negative interactions among these elements (Figure 2 B and C).

**Figure 2.**
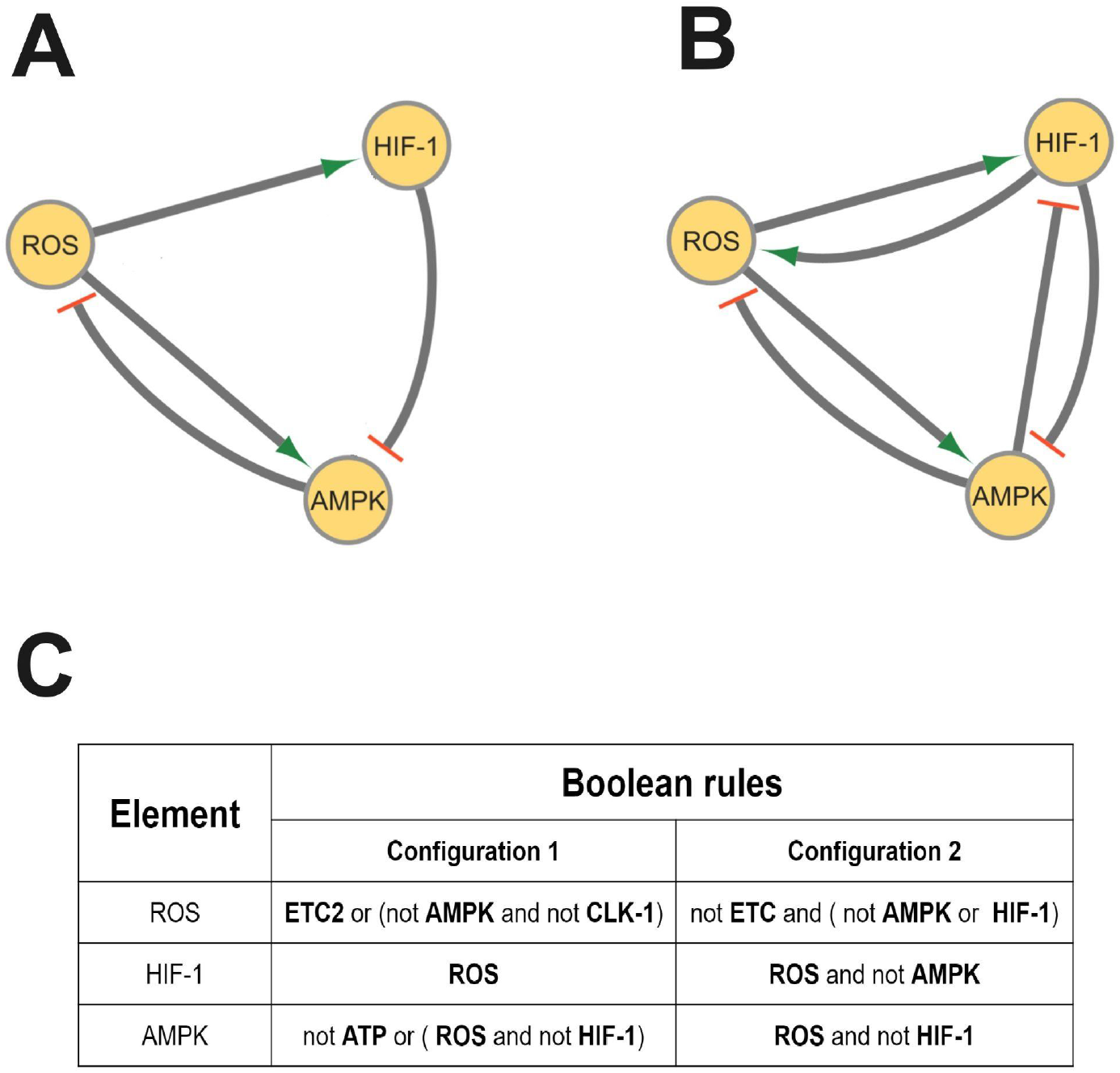
The dual-configuration for the ROS, HIF-1 and AMPK loop. For a better depiction of the system in a discrete model it was necessary to establish a new and simpler configuration (A). Graph of the configuration 1 (the simpler configuration) for the *clk-1* network (B). Graph of the configuration 2 (the accurate configuration) for the *clk-1* network (C). Boolean rules for both configurations.

This dual-configuration strategy underscores our commitment to accurately capturing the nuances of the *clk-1* regulatory network, acknowledging the need for flexibility in modeling approaches to accommodate the inherent complexity of biological systems. Through this detailed examination of the *clk-1* interactions and the strategic implementation of modeling techniques, we aim to shed light on the underlying regulatory mechanisms contributing to the *clk-1* mutant’s unique phenotype, offering valuable insights into the broader context of mitochondrial function and its implications for cellular aging.

### The long lived mutant attractors show upregulation in all processes

The steady states of the model are called attractors and, in a discrete approach, these attractors are the states, or combinations of states, where all the other possible states are led. For configuration 1, in the wild type (N2 strain) conditions, the value of *clk-1* is always 1 (active) and 100% of the states drive to a single state attractor where all the main processes are inhibited (Figure 3A), but in mutant conditions when the *clk-1* value is always 0 (inactive) the main attractor is composed of 16 states where all the processes are oscillating between active and inactive (Figure 3B), also a 78% of all possible states led to this attractor. Besides this result, two more cyclic attractors of 11 and 10 states exist in the *clk-1* mutant conditions (Figure S1). These attractors show the same result as the 16-state attractor, where all the processes oscillate.

**Figure 3.**
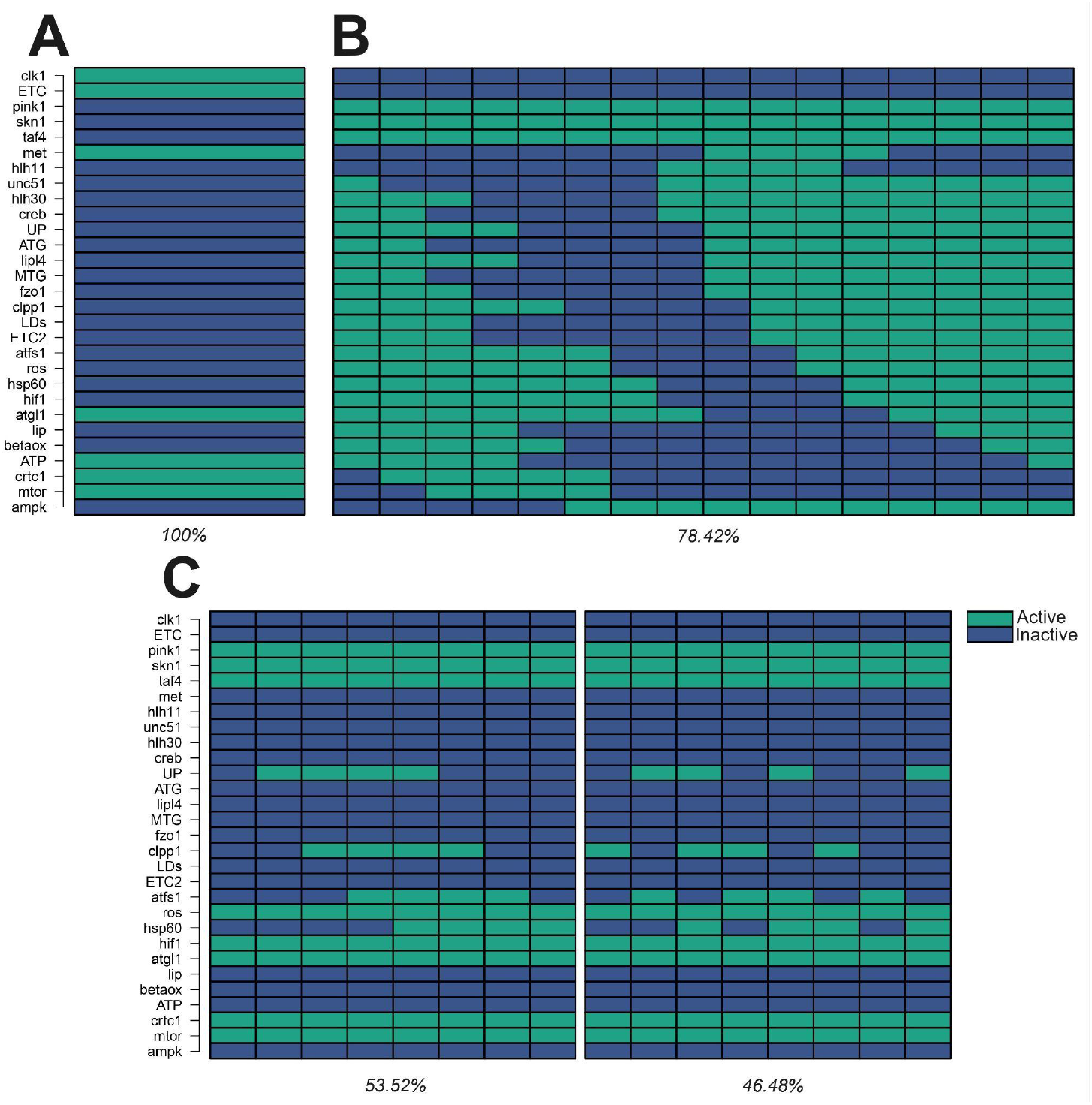
The logical modeling analysis reflects the impact of molecular interactions on mutant strains. Computation of the attractors for all nodes of the model in three different conditions: columns are the states and lines the nodes, A color code denotes each node state, active (green) or inactive (blue) at the bottom of each attractor is shown its basin of attraction (A). Static attractor for the wild type N2 condition (B). Main cyclic attractor for the *clk-1* mutant condition (C). The two cyclic attractors for the double mutant *clk-1;aak-2* condition.

The other condition tested was the *clk-1;aak-2* double mutant, where the *clk-1* and AMPK value is always 0; the two resulting attractors show a static condition similar to the wild type attractor, unlike this condition, the elements associated with the UPRmt are oscillating. Also, the elements *pink-1* and *skn-1* are activated (Figure 3C), and both attractors have a basin of attraction close to 50%. These results so far were obtained for configuration 1 in a discrete approach, the major difference within the first and second configuration pivot on the *clk-1* mutant conditions. While configuration 1 throws feasible attractors, configuration 2 throws a set of 10 cyclic attractors, and none of those cyclic attractors contemplates the electric transport chain compensation (Figure S2). In addition to this condition, both configurations’ attractors for the double mutant and the wild-type conditions are the same.

The discrete attractors for configuration 1 showed a better simulation of the *clk-1* mutant conditions. We found that the inductor of UPRmt, the *atfs-1* transcription factor, increases its expression in this mutant (27); for the autophagy process, the *hlh-30* transcription factor increases its expression (8), in the case of mitochondrial fusion there is an increase in the number of elongated mitochondria as well as an expression increase of the mitofusins *eat-3* and *fzo-1* (28), finally a microarray analysis revealed that the genes involved in fatty acid oxidation increase their expression (29).

### The continuous approach is a better representation of the system

In the continuous approach, the logical rules are fitted into ordinary differential equations with sigmoidal functions; each element can constantly change its activation state according to the combination of positive and negative interactions. For configuration 1, the continuous attractor shows a constant oscillation for each element (except for the static elements in the discrete approach) (Figure 4A); some groups of elements in this configuration show an oscillation pattern where these groups are activated and inhibited at the same time (Figure S3). On the other hand, configuration 2 shows a more feasible result than the discrete approach, in this configuration, the elements reach a steady state on a static value between 0 and 1, but in most of the cases is not 0 or 1 (Figure 4B and S2B). This result could be an artifact of the model; to prove this, further research is required. While configuration 2 might reflect the general expression level, configuration 1 may reflect the expression dynamics at a single cell level. These stable states are particularly interesting as they represent the network configurations that can persist over time, potentially correlating with specific phenotypes.

**Figure 4.**
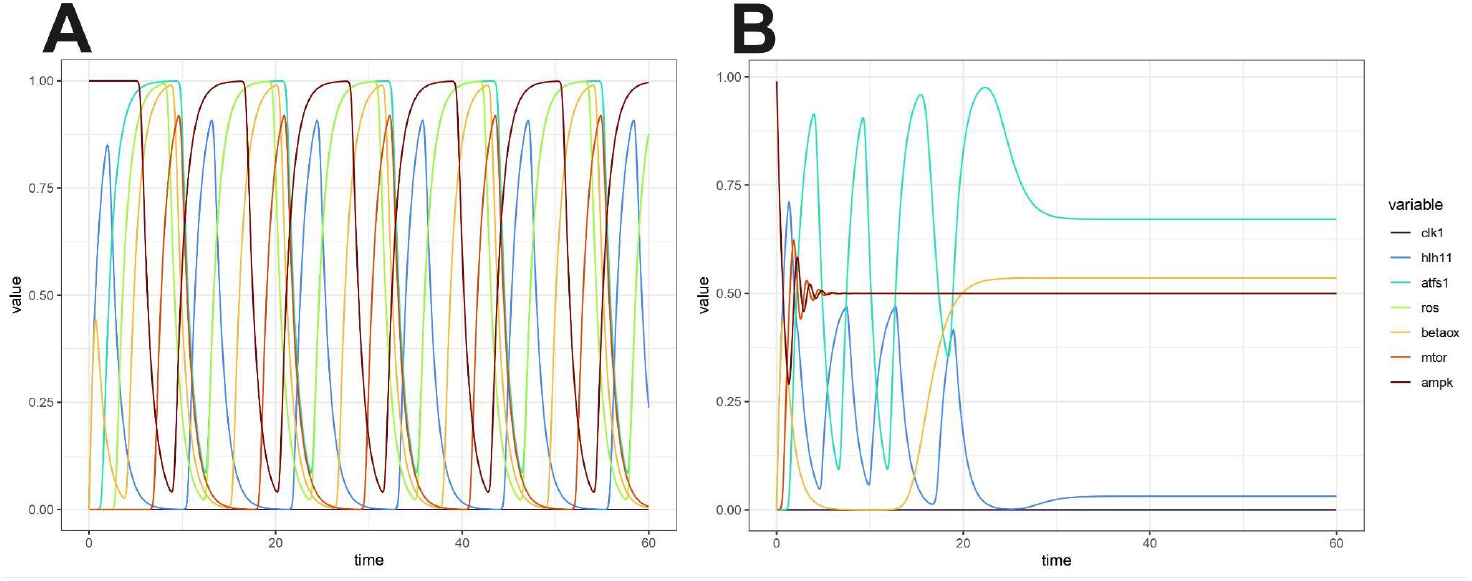
Configuration 2 has a static attractor for the *clk-1* mutant condition in the continuous approach. Computation of the stable states on selected nodes of the model in the *clk-1* mutant condition, each line depicts a node. The Y axis shows the activation level between 0 (inactive), and 1 (active) and the × axis shows the time. (A) The continuous simulation for the configuration 1 cyclic attractor (B) The continuous simulation for the configuration 2 static attractor.

### The healthspan is unrelated to lifespan

The lifespan was measured through a survival assay, we found that N2, *aak-2* and *clk-1;aak-2* have a shorter lifespan than *clk-1* worms (Figure 5), a Cox regression analysis revealed that there is a significative difference between N2 and the strains *aak-2(ok524)* and *clk-1(qm30)*, but not with the strain *clk-1(qm30);aak-2(ok524)*, also there is a 71.5% less risk of death for the *clk-1* concerning the strain N2.

**Figure 5.**
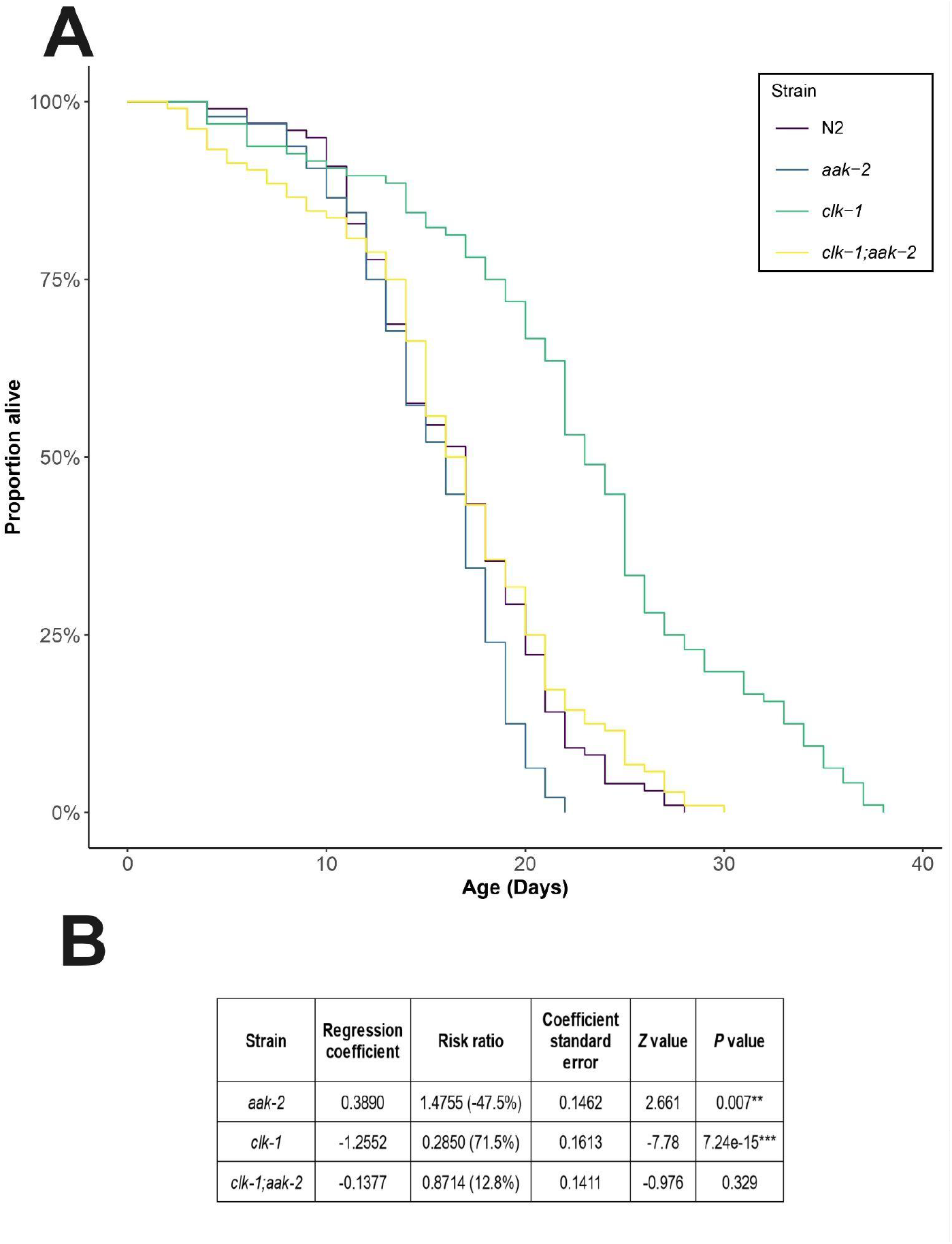
Only the *clk-1* strain has a positive effect on lifespan. (A) Survival curve of all strains used in this study: wild type (N2) (purple), *aak-2(ok524)* (blue), *clk-1(qm30)* (green), and *clk-1(qm30);aak-2(ok524)* (yellow). (B) Cox regression analysis based on the hazard ratio of the mutant strains compared with the wild type. The significance codes are the following: P = 0, ‘***’; P < 0.001, ‘**’; P < 0.01, ‘*’ and, “ “, non-significant.

We proposed a novel estimation of a health index based on the general muscular attrition in *C. elegans* to measure its health condition. This variable is measured in 1-day adult worms until they reach the 9-day age and is through the behaviors of pharyngeal pumping, swimming, and defecation. We found significant differences between strains and measurement times for individual and aggregated behaviors (Figure S4).

We implemented two methods to estimate the health index. The first one is based on symbolic regression estimation. This method uses a genetic algorithm to calculate the best function that fits the data, and then we take the definite integral of this function to measure the area under the curve. Below is shown an example of this kind of calculation using the N2 data estimated function.

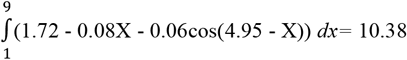

The second method is a direct measurement from the data; we defined a line through the points established by the quartiles of each day’s measurement, and then we calculated the area of the polygons defined by this line and the distance between measurements, as shown in Figure S5. The area under the function, or the quartiles, reflects the general attrition. If the behavior keeps high and healthy longer, the area (index) will be bigger, but if the behavior is slow and decays quickly, the index will be smaller.

The results show that strain N2 has a higher health index (therefore the best health condition), and strain *aak-2(ok524)* is close to this value. On the other hand, the long-lived strain *clk-1 (qm30)* has almost half of this value, and the double mutant *clk-1(qm30);aak-2(ok524)* is closer to a quarter of the N2 value (Table 2). The general result is replicated in any aggregate of behaviors as well as in individual behaviors (Table S1), despite this information, the more behaviors measured are aggregated to the health index, the bigger the impact shown on ill strains, and the health condition of the strains is not as positive as their lifespan. We also checked which strains had the greatest deterioration according to their initial conditions; the result between strains was the same, but there were slight differences (Table S2). Regarding the two methods used, the quartile 2 calculated area fits the symbolic regression estimated area (Tables 2, S1, and S2), this makes the area calculation method more accurate than the symbolic regression method but makes the regression method more useful for longitudinal studies where the data is a cloud of points.

**Table 2.** Health index estimation. The area from the symbolic regression method was calculated through the definite integral of each estimated function, and the area from the quartiles was calculated from the polygons drawn by the data quartiles of each day.

| Strain | Symbolic regression estimation |  |  | Area under the quartiles |  |  |
| --- | --- | --- | --- | --- | --- | --- |
|  | Regression function | Area | % | Q1 | Q2 | Q3 |
| **Pharyngeal pumping and swimming index** |  |  |  |  |  |  |
| N2 | $Y = 1.65 + 0.05\sin(6.03 + X) - 0.06X$ | 10.91 | 100% | 10.028 | 10.896 | 11.639 |
| *aak-2* | $Y = 1.73 + 0.01/\sin(X) - 0.08X$ | 10.55 | 96.7% | 10.12 | 10.699 | 11.469 |
| *clk-1* | $Y = 1.28 + 4.62e^{-11}X(X^X) - 0.11X$ | 5.84 | 53.52% | 5.139 | 5.966 | 6.95 |
| *clk-1;aak-2* | $Y = 97.2/(33.57 + X) - 2.12$ | 3.25 | 29.78% | 2.44 | 3.263 | 3.971 |
| **Pharyngeal pumping, swimming and defecation index** |  |  |  |  |  |  |
| N2 | $Y = 1.72 - 0.08X - 0.06\cos(4.95 - X)$ | 10.38 | 100% | 9.299 | 10.279 | 10.896 |
| *aak-2* | $Y = 1.83 + 0.06\cos(1.8X) - 0.11X$ | 9.94 | 95.76% | 9.416 | 10.03 | 10.662 |
| *clk-1* | $Y = 0.57 + 0.63\sin(X^2)$ | 4.8 | 46.24% | 2.018 | 3.105 | 4.179 |
| *clk-1;aak-2* | $Y = 0.71 - 0.1X - 6.32e^{-6}X^5$ | 1.06 | 10.21% | 1.214 | 1.738 | 2.249 |

## Discussion

Until now, most research on the aging process has focused on single gene interactions, ignoring the underlying molecular network or the health condition and lifespan status in the case of computational modeling. Our work aimed to address these limitations by first inferring the molecular network involved in the *clk-1* aging phenotype and then testing the Boolean model to identify steady-state conditions for relevant genes in this mutant. Subsequently, using bioassays and a novel health index estimation, we established the lifespan and health conditions of relevant mutants tested in the Boolean model. This comprehensive approach enabled us to elucidate the regulatory dynamics involved in the aging process.

This study showed that under certain conditions, such as those involving the mitochondrial mutant *clk-1*, various processes are triggered, indicating a stress response. This stress response was wholly or partially inhibited in the wild type and double mutants of *the clk-1* and *aak-2* strains, respectively (Figures 3A and 3C). Under these conditions, both strains had similar lifespans (Figure 5), implying that the stress response is involved in the *clk-1* mutant lifespan extension (Figures 3B and 5). On the other hand, health index results showed a decreased health condition for the *clk-1*mutant (Table 2), contradicting hormesis theory (9). However, adverse conditions still have a beneficial effect on lifespan.

In contrast, the double mutant *clk-1;aak-2*, did not have a long lifespan or good health span (Figure 5 and Table 2), which can be attributed to the absence of cell repair and increased levels of reactive oxygen species. These findings indicate that the lifespan of *C. elegans* is dependent on its stress response and subsequent metabolic, epigenetic, and cellular changes. However, health span is only affected by the amount of cellular stress and the extent to which it is managed.

Estimating a health index helps measure the health conditions in this study. It does not assume a specific shift from healthy to ill, thus, providing a more comprehensive view of health. This index reflects the development of sarcopenia, which is influenced by muscular attrition and behavior. It can occur in the *clk-1* mutant strain, despite its initial slow rate of behavior, which does not slow down metabolism and instead increases the lifespan (30). Therefore, slow behavior is linked to the development of sarcopenia in these strains.

Regarding the model, one noteworthy aspect of the attractors obtained was their cyclic properties, which may approximate the theory of homeodynamics (31). This theory suggests that biological systems are not static, and should be studied as complex processes with emerging properties. Aging is a complex process due to its hierarchical structure and how a healthy or frail condition emerges as an emergent property of molecular and cellular dynamic interactions (32). Under this assumption, cyclic attractors can be interpreted as a form of dynamic equilibrium from which lifespan and health condition phenotypes emerge. For this model, defining the dynamic balance that regulates the conditions of *clk-1*, including all metabolic, epigenetic, and proteostatic factors, and determining whether the N2 attractor remains static with appropriate information and how a cyclic attractor changes over time is necessary.

Analyzing the aging process as an emergent property of homeodynamics might bring us closer to defining an aging paradigm, as the hallmarks of aging have been identified thus far (33). Understanding the role of each molecular factor and its contribution at the primary or secondary level is essential for defining the aging paradigm (34). Logical modeling is an excellent tool for integrating and analyzing these factors as a dynamic and complex process. Nonetheless, it is crucial to demonstrate that gene expression correlates with cyclic attractors and follows a homeodynamic pattern. To achieve this, it is necessary to measure genetic expression at the cellular level.

## Conclusions

In our study, we used logical modeling and quantitative methods to investigate the relationship between the processes associated with the *clk-1* mutant and its aging phenotypes. This comprehensive approach enabled us to elucidate the potential homeostatic properties of the aging process, highlighting the critical role of stress responses and dynamic equilibrium as key regulators of lifespan. Our findings indicate promising directions for future exploration of this domain.

Estimating health conditions using a health index provides a comprehensive view, reflecting the development of sarcopenia influenced by muscular attrition and behavior. The cyclic properties of the attractors in the model suggest a homeodynamic theory, which posits that homeostasis is not a static process, as the name implies, but a dynamic system in which all metabolites constantly change in concentration and interaction and how these changes evolve may explain the aging process

Analyzing aging as an emergent property of homeodynamics may help define an aging paradigm, as identifying primary and secondary aging causes remains challenging owing to the scale of the aforementioned dynamics and the complex system properties of this process. Logical modeling is valuable for integrating and analyzing molecular factors in dynamic processes. However, demonstrating the correlation between gene expression, cyclic attractors, and homeodynamic patterns requires precise cellular-level measurements using advanced technologies.

## Acknowledgments

We thank FONDEC-UAQ 2022 (20205007011001) and for their financial support. Additionally, we acknowledge CONAHCyT for providing a Master’s degree stipend (805150).

