## Supplementary material for "Stress responses and dynamic equilibrium: Key determinants of aging in the *C. elegans clk-1* mutant": Figure S1, Figure S2, Figure S3, Figure S4, Figure S5, Table S1, Table S2

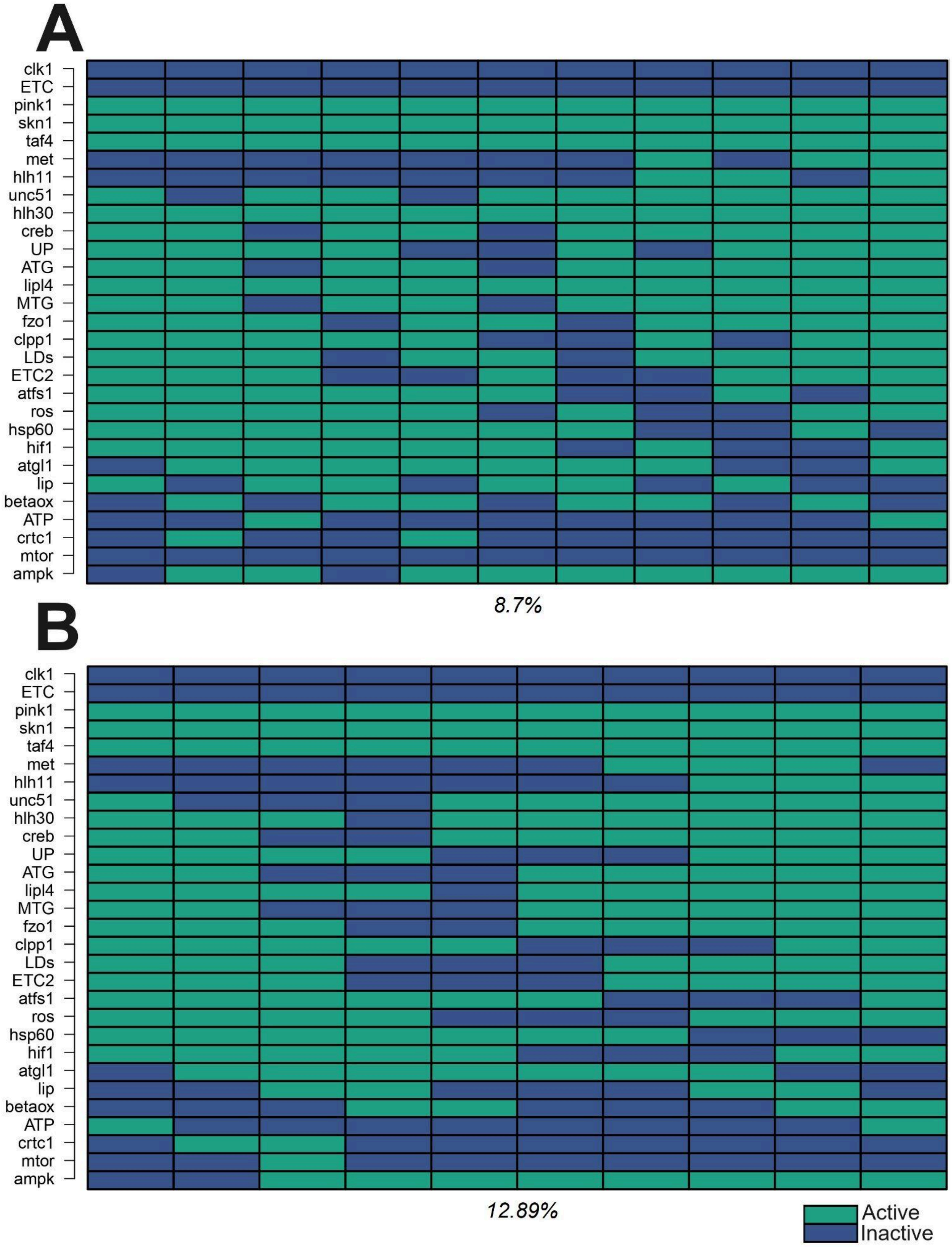


**Figure S1. The *clk-1* secondary attractors.** Computation of the attractors for all nodes of the model for the *clk-1* mutant condition, columns are the states and lines the nodes, A color code denotes each node state, active (green) or inactive (blue) at the bottom of each attractor is shown its basin of attraction (A). The 11 state attractor with the smallest basin of attraction .(B) The 10 state attractor.

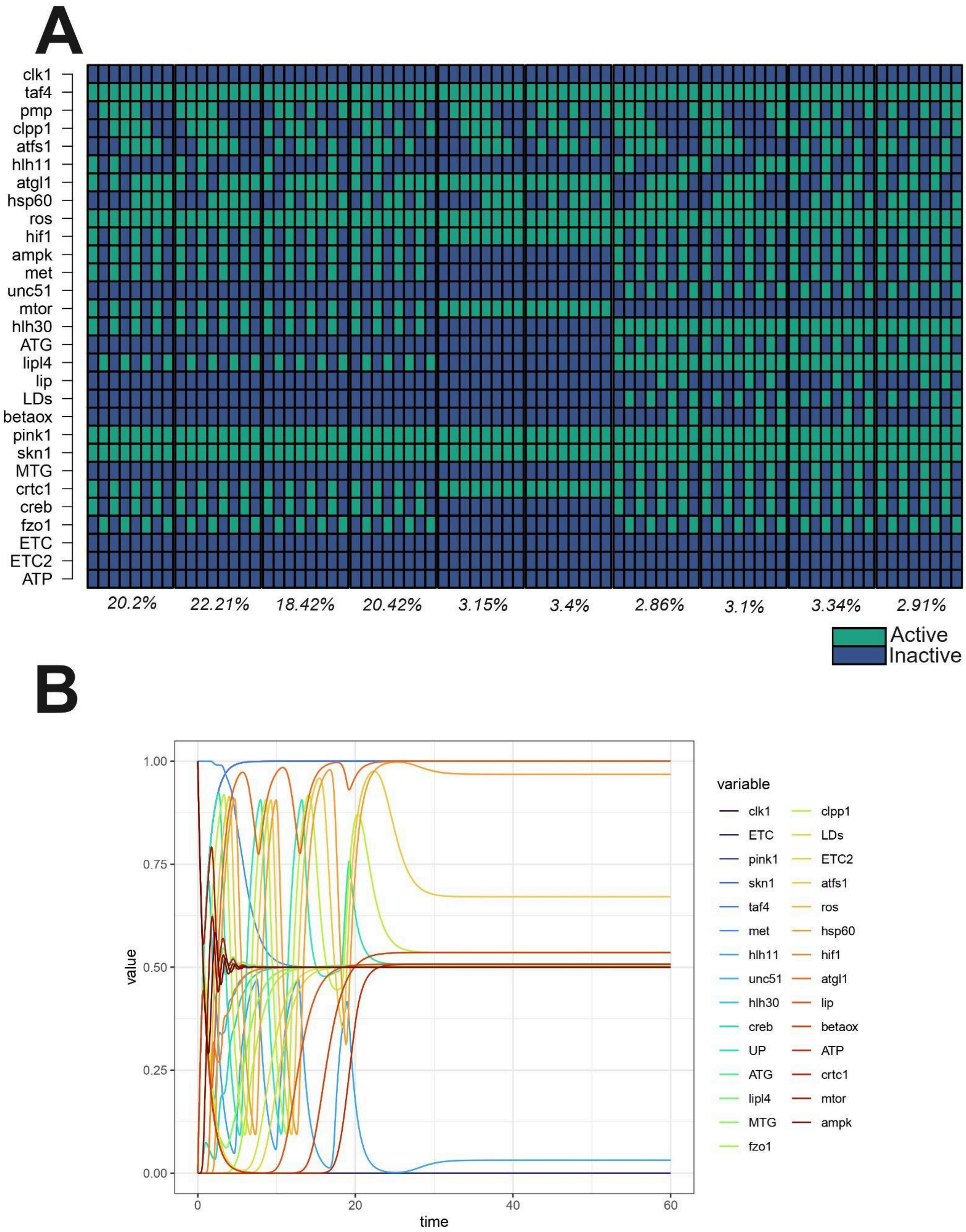


**Figure S2. The discrete and continuous attractors for the configuration 2 in the *clk-1* mutant conditions.** Computation of the stable states for the configuration 2 in the *clk-1* mutant condition. (A) 10 discrete attractors of 8 states each (B) The ecstatic attractor for the continuous approach.

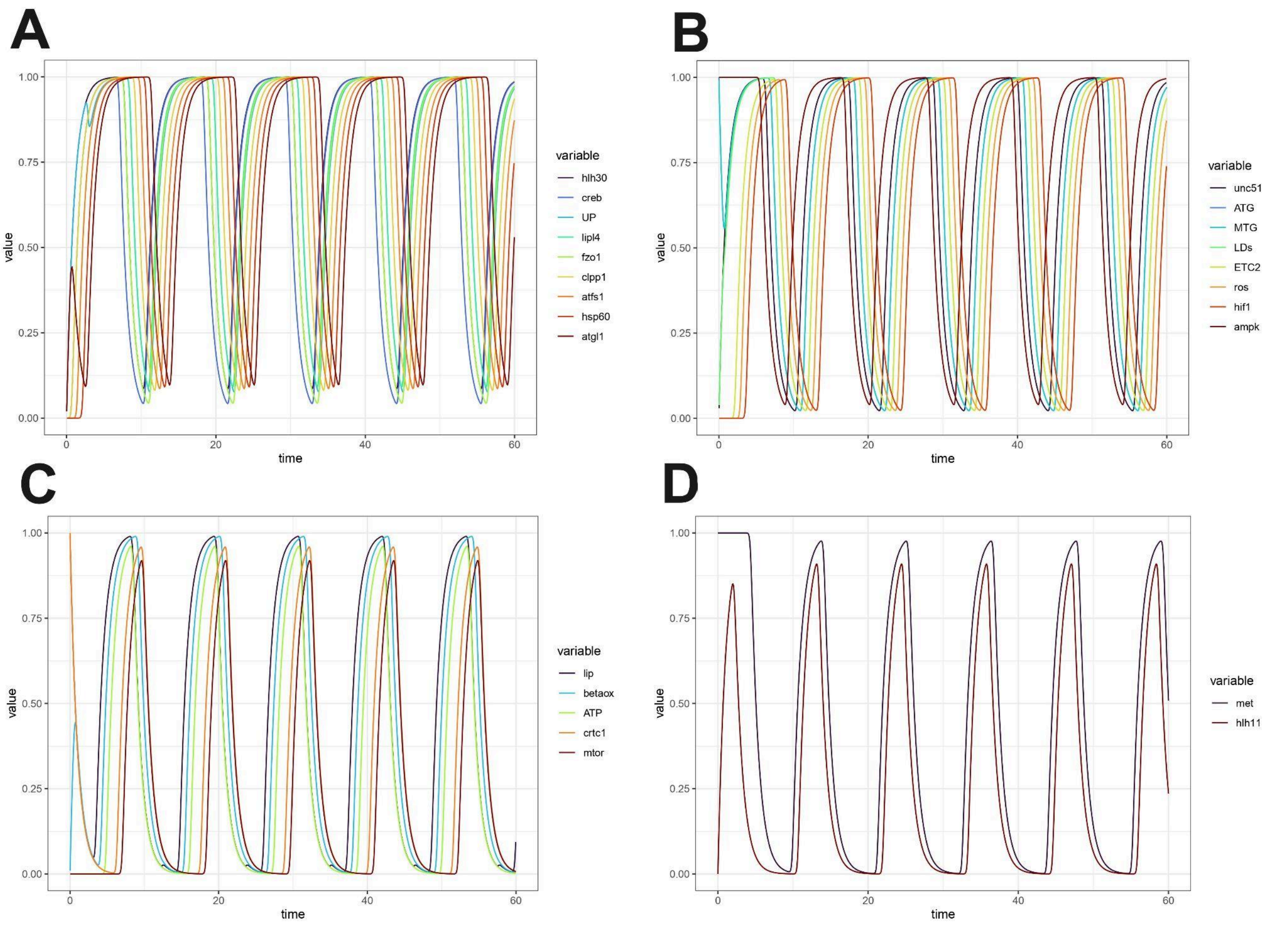


**Figure S3. The configuration 1 patterns of oscillation for the *clk-1* mutant conditions.** Computation of the stable states on selected nodes of the model that share a pattern of oscillation in the *clk-1* mutant condition, each line depicts a node. The Y axis shows the activation level between 0 (inactive) and 1 (active) and the X axis shows the time. (A) pattern 1(B) pattern 2 (C) pattern 3 (D) pattern 4.

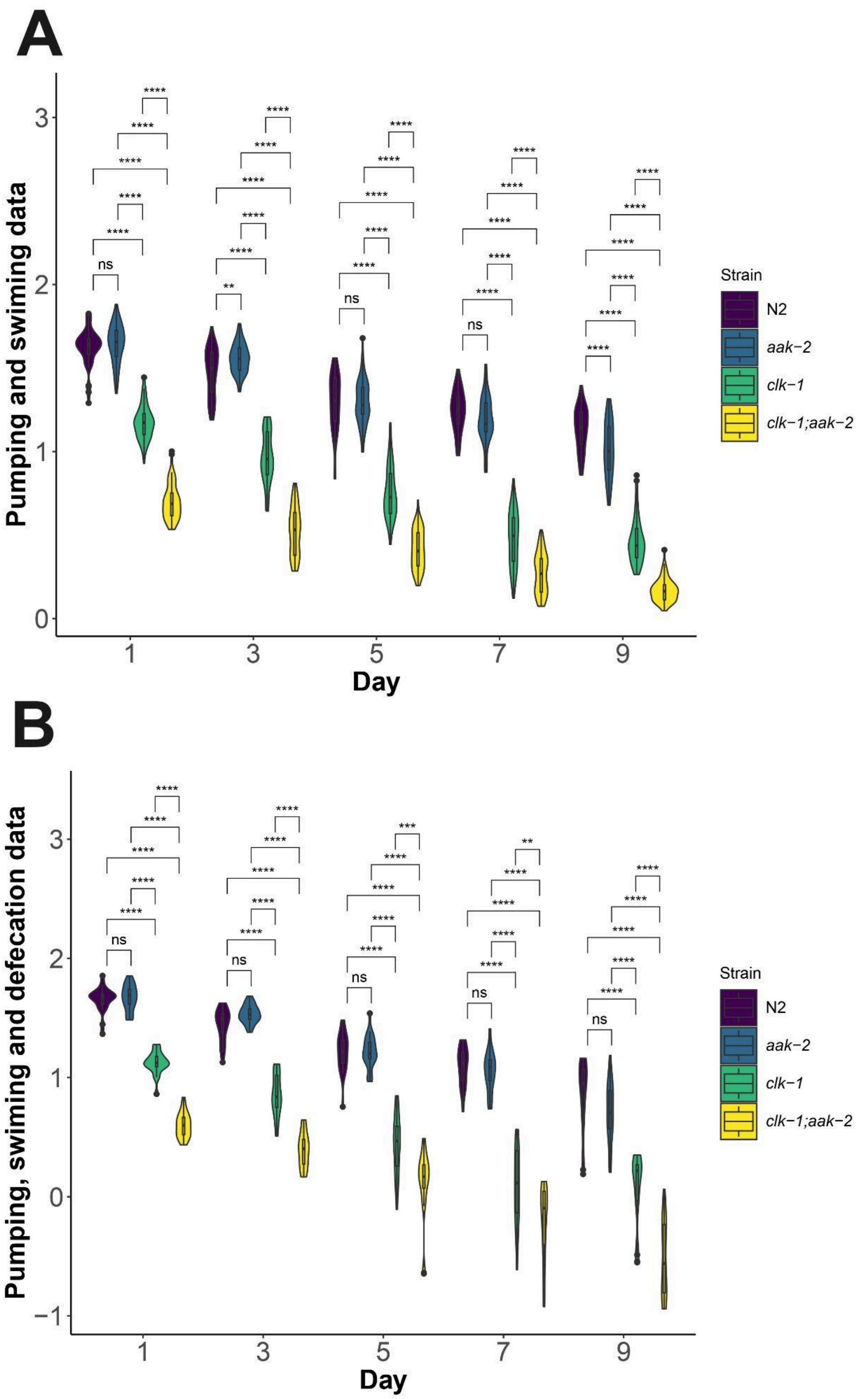


**Figure S4. The neuromuscular behaviors data**. (N2) (purple), *aak-2(ok524)* (blue), *clk-1(qm30)* (green) and *clk-1(qm30);aak-2(ok524)* (yellow) (A) Aggregated data for the pharyngeal pumping and swimming behaviors. (B) Aggregated data for the pharyngeal pumping, swimming and defecation behaviors. The two way mixed ANOVA significance codes are the following: P = 0, '***'; P < 0.001, '**'; P < 0.01, '*' and, " ns", non significant.

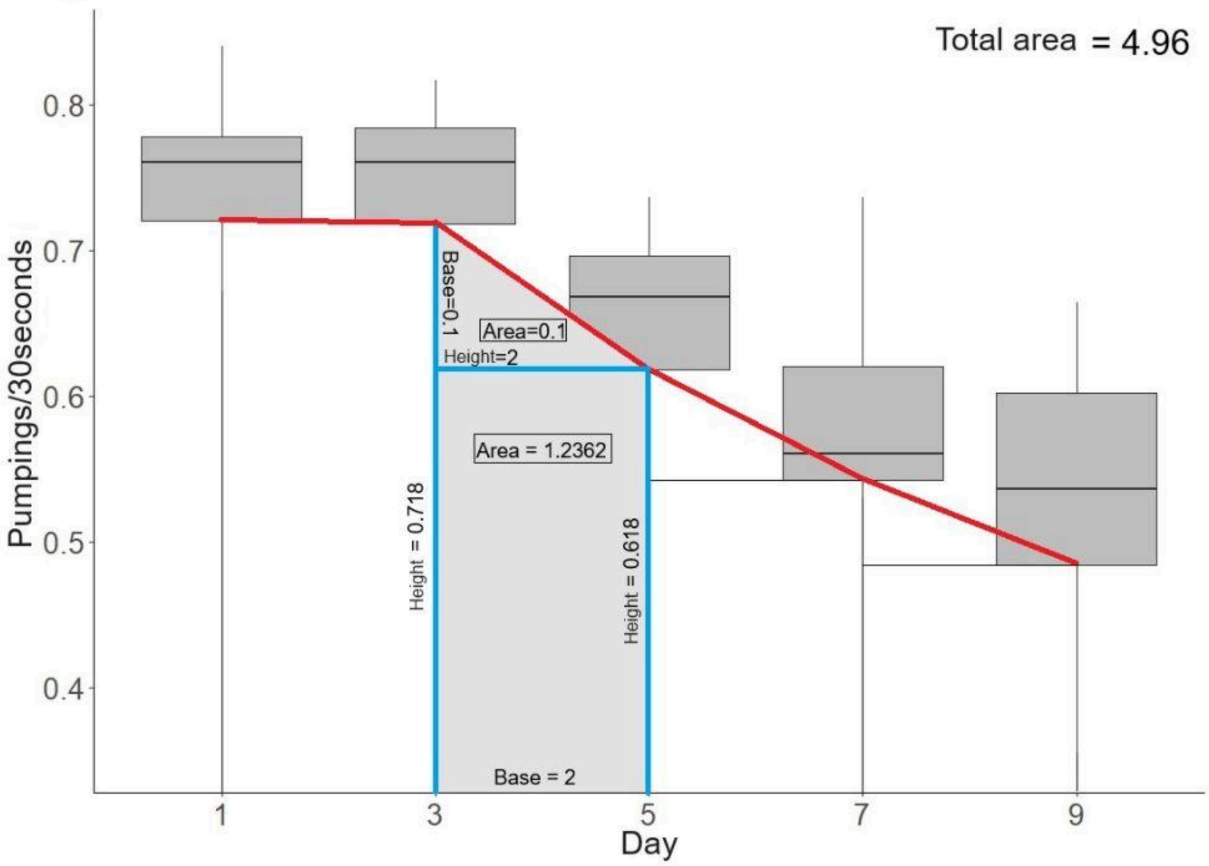


**Figure S5. Health index estimation**. In the Direct Area Calculation from Behavioral Data, the red line shows under which quartile the area is calculated. The blue lines delimit the polygons, they are defined by the quartile value in the red line, the differences in the height between quartiles, and the distances between measurements. finally the area of each polygon is calculated and summed.

**Table S1. Health index estimation for individual behaviors.** The area from the symbolic regression method was calculated through the definite integral of each estimated function, and the area from the quartiles was calculated from the polygons drawn by the data quartiles of each day.

| Strain | Symbolic regression estimation | | | Area under the quartiles | | |
|---|---|---|---|---|---|---|
| | Regression function | Area | % | Q1 | Q2 | Q3 |
| **Pharyngeal pumping behavior index** | | | | | | |
| N2 | $Y = 0.94 + 0.03\sin(X) - 0.05X$ | 5.19 | 100% | 4,414 | 5,162 | 5,654 |
| *aak-2* | $Y = 0.90 + 0.005\sin(X) - 0.04X$ | 5.32 | 102.5% | 4,987 | 5,329 | 5,608 |
| *clk-1* | $Y = 0.42 - 0.2\sin(4.09 - 5.78X)$ | 3.42 | 64.28% | 2,608 | 3,237 | 3,850 |
| *clk-1;aak-2* | $Y = -10.96/(-34.11 - X^2)$ | 1.54 | 29.67% | 1,035 | 1,559 | 2,418 |
| **Swimming behavior index** | | | | | | |
| N2 | $Y = 0.72 + 0.05\sin(2.9828^X)$ | 5.75 | 100% | 5,349 | 5,906 | 6,260 |
| *aak-2* | $Y = 0.86 - 0.03X - 0.03\cos(0.22 - 1.14X)$ | 5.47 | 95.13% | 4,806 | 5,436 | 6,113 |
| *clk-1* | $Y = 0.57 + 0.01\cos(0.48 + 1.05X) - 0.04X$ | 2.8 | 48.69% | 2,309 | 2,813 | 3,308 |
| *clk-1;aak-2* | $Y = 0.36 + 0.03\sin(284.65X^2) - \sin(0.03X + 0.001\sin(0.36X))$ | 1.57 | 27.3% | 1,186 | 1,452 | 1,787 |
| **Defecation behavior index** | | | | | | |
| N2 | $Y = 0.21 + -0.95/(3.68 + X)$ | 0.79 | 100% | 0.652 | 0.788 | 1.057 |
| *aak-2* | $Y = X/(56.56 + 1.29X\cos(X) - X)$ | 0.79 | 100% | 0.689 | 0.808 | 1.036 |
| *clk-1* | $Y = 0.03X + 0.03\cos(X)$ | 1.56 | 197.4% | 1.036 | 1.076 | 2.637 |
| *clk-1;aak-2* | $Y = 0.08 + 0.0008X^3$ | 2.02 | 255.6% | 1.467 | 2.186 | 3.472 |

**Table S2. Health index estimation for individual strains.** The data were changed into relative scales for each strain to explore the general decay of each strain with respect to its initial conditions; the percentage shows the value reached with respect to the maximum value where there is no deterioration.

| Strain | Symbolic regression estimation | | | Area under the quartiles | | |
|---|---|---|---|---|---|---|
| | Regression function | Area | % | Q1 | Q2 | Q3 |
| **Pharyngeal pumping behavior index** | | | | | | |
| N2 | $Y = 0.93 + 0.04\sin(0.97X) - 0.07X$ | 4.61 | 51.22% | 3,697 | 4,594 | 5,185 |
| *aak-2* | $Y = 0.92 + 0.009\sin(5.97 + X) - 0.06X$ | 4.94 | 54.88 | 4,499 | 4,942 | 5,306 |
| *clk-1* | $Y = 0.85 - 0.07X - 0.01X\cos(0.01 +X)$ | 3.75 | 41.66% | 2,967 | 3,76 | 4,532 |
| *clk-1;aak-2* | $Y = 0.72 - \sin(0.06 + 0.05X)$ | 3.14 | 34.88% | 2,038 | 3,07 | 4,761 |
| **Swimming behavior index** | | | | | | |
| N2 | $Y = 0.65 + 0.01X\cos(X) - 0.14\cos(0.47 + X)$ | 5.4 | 60% | 4,253 | 5,349 | 6,045 |
| *aak-2* | $Y = 0.68 - 0.005X^2 - 0.07\cos(1.15 + 0.92X)$ | 4.28 | 47.55% | 3,290 | 4,218 | 5,218 |
| *clk-1* | $Y = 0.77 - 0.06X - 0.01\sin(0.98 + 2.09X)$ | 3.69 | 41% | 2,309 | 2,813 | 3,309 |
| *clk-1;aak-2* | $Y = 0.7 + 0.06\cos(X) + 0.0001X^4 - 0.05X - 0.001X^3$ | 1.86 | 20.66% | 2,257 | 2,764 | 3,402 |
| **Defecation behavior index** | | | | | | |
| N2 | $Y= 0.15X - 0.08 - 0.009X^2$ | 3.05 | - | 2.452 | 3.8 | 3.479 |
| *aak-2* | $Y = 0.74X/(10.9810 + \cos(X)) - 0.03$ | 2.41 | - | 2.053 | 2.417 | 2.873 |
| *clk-1* | $Y = 0.03X + 0.03 \log(x)\cos(5.96 + X)$ | 1.41 | - | 0.575 | 1.337 | 2.45 |
| *clk-1;aak-2* | $Y = 0.0078X^2$ | 1.89 | - | 1.206 | 1.973 | 3.305 |
| **Pharyngeal pumping and swimming index** | | | | | | |
| N2 | $Y = 1.61 + 0.01X\sin(X) - 0.08X$ | 9.77 | 54.27% | 8,403 | 9,744 | 10,857 |
| *aak-2* | $Y = 1.72 - 0.11X - 0.06\cos(5.13X)$ | 9.04 | 50.22% | 8,23 | 9,053 | 10,212 |
| *clk-1* | $Y = 1.61 - 0.14X - 2.4e^{-5}\cos(X)\exp(X)$ | 7.23 | 40.16% | 6,304 | 7,32 | 8,602 |
| *clk-1;aak-2* | $Y = 1.48 + 0.003X^2 - 0.15X$ | 6.28 | 34.88% | 4,714 | 6,323 | 7,713 |
| **Pharyngeal pumping, swimming and defecation index** | | | | | | |
| N2 | $Y = 0.32 + 1.73/(X + \cos(1.15X))$ | 7 | 38.88% | 5,42 | 6,949 | 8,084 |
| *aak-2* | $Y = 1.66 + 0.16\sin(6.1 - 49.39X) - 0.16X$ | 6.78 | 37.66% | 5,786 | 6,852 | 6,936 |
| *clk-1* | $Y = 1.66 + 0.15\cos(X) - 0.18X - 0.05X\cos(X)$ | 5.62 | 31.22% | 4,507 | 5,815 | 7,458 |

| | | | | | | |
|---|---|---|---|---|---|---|
| *clk-1;aak-2* | $Y = 1.51 - 0.18X - 3.34e^{-6}X\exp(X)$ | 4.62 | 25.66% | 3,429 | 4,797 | 5,928 |
